# Incorporating genetic selection into individual-based models (IBMs) of malaria and other infectious diseases

**DOI:** 10.1101/819367

**Authors:** Ian M. Hastings, Raman Sharma

## Abstract

Optimal control strategies for human infections are often investigated by computational approaches using individual-based models (IBMs). These typically track humans and evaluate the impact of control interventions in terms of human deaths, clinical cases averted, interruption of transmission etc. Genetic selection can be incorporated into these IBMs and used to track the spread of mutations whose origin and spread are often driven by the intervention, and which subsequently undermine the control strategy; for example, mutations which encode antimicrobial drug resistance or diagnosis- or vaccine-escape phenotypes. Basic population genetic descriptions of selection are based on infinite population sizes (so that chance fluctuations in allele frequency are absent) but IBMs track finite population sizes. We describe how the finite sizes of IBMs affect simulating the dynamics of genetic selection and how best to incorporate genetic selection into these models. We use the OpenMalaria IBM of malaria as an example, but the same principles apply to IBMs of other diseases. We identify four strategies to incorporate selection into IBMs and make the following four recommendations. Firstly, calculate and report the selection coefficients, s, of the advantageous allele as the key genetic parameter. Secondly, use these values of ‘s’ to calculate the wait-time until a mutation successful establishes itself in the population. The wait time for the mutation can be added to speed of selection, s, to calculate when the mutation will reach significant, operationally important levels. Thirdly, quantify the ability of the IBM to robustly estimate small selection coefficients. Fourthly, optimise computational efficacy: when ‘s’ is small it is plausible that fewer replicates of larger IBMs will be more efficient than a larger number of replicates of smaller size.

## 1. Introduction

Advances in computational power over the last 20 years have allowed sophisticated, individual-based models (IBMs) of infectious diseases to be developed and applied to important human and animal disease. These are particularly valuable in diseases with complex transmission through vector species and/or complex clinical aetiology because they allow considerably more realism to be added to the fundamental susceptible-infected-recovered (SIR) models which are usually the first analyses used to investigate disease epidemiology. The next challenge for these IBMs is to incorporate genetic selection into the disease epidemiology. Most pathogens readily evolve in response to human interventions. For example, drug resistance almost inevitably evolves in response to drug deployment, and mutations arise that change the antigenic profile of the molecules detected by molecular diagnosis, making the pathogens invisible to diagnosis. Incorporating this evolution into IBMs is not straightforward and this manuscript discusses how it may best be achieved in terms of accurately quantifying the selection process in a computationally efficient manner.

Our expertise lies in malaria and IBMs of malaria have been recently, and comprehensively, reviewed by Smith and colleagues [1]. The development of several IBM models was supported by the Bill and Melinda Gates Foundations and the resulting consensus exercises co-ordinated by these groups has been influential in evaluating the potential impact of intervention such as partially-effective vaccines [2], mass drug administration (MDA) programmes [3], and the impact of new diagnostic tools [4]. These publications focused on the relatively short-term impact of interventions but the historic problem with such interventions is that the malaria populations evolve and adapt to the challenges posed by the interventions. The obvious examples are vaccine-based interventions driving vaccine-insensitive variants (e.g. [5, 6] and discussion in [7]), drug-based interventions driving drug resistance (e.g. Figure 3 of [8]), and deployment of rapid diagnostic test (RDTs) driving mutations that prevent the infection being detected and diagnosed (such as *hrp2*- deletion mutants) [9, 10]. The basic dynamics of spread of such mutations can be obtained from standard population genetics methodology (e.g. [11–13] and later). However, the simplicity of these methodologies means the analyses must focus on malaria genetics and often largely ignore the epidemiological and clinical frameworks within which these selection processes operate. Consequently, there is increasing interest in incorporating genetic selection into the IBMs of malaria. The use of IBM makes the selective background far more realistic and can incorporate factors such as heterogeneity in mosquito biting, the effectiveness of diagnosis, the effect of human acquired immunity, the impact of super-infection, local patterns of clinical treatment, amongst other factors.

There is a considerable amount of population genetic theory that many epidemiological modellers may be unaware of, presumably through their background as mathematicians and/or computer scientists. Population genetic theory developed over the last 100 years largely in the absence of computer infrastructure so there is a large amount of basic theory that can be combined with large-scale IBMs of disease transmission to make selection more transparent, more computationally efficient and, importantly, to make the results and outputs comparable across simulation platforms. The aim of this paper is to describe how this may be achieved. The significant differences between the two approaches is that simple populations genetic theory usually assumes infinite population sizes (so that random fluctuations in allele frequency are absent) whereas IBMs track finite population sizes. There are two related effects that occur in finite populations that affect how we bring genetic selection into IBMs and these need to be understood before we describe the integration of population genetics into IBMs. Both effects arise because IBMs track numbers, rather than frequencies, of each genotype of parasites:

> *Stochastic fluctuation may result in “genetic extinction” when small numbers of one allele type are present*. This effect emphasises the difference between the expected change in the number of infections carrying the allele, and the actual change. Suppose there are 10 infections carrying the advantageous allele (e.g. drug resistance) in the IBM and the selection coefficient (see later) acting on the allele is 0.05, i.e. the allele frequency is expected to increase by 5% per malaria generation which is roughly in line with field estimates (see later). The expected number of resistant infections next generation is 10*1.05= 10.5 but it is obviously impossible to leave exactly 10.5 infections: the number of resistant infections must come from a distribution i.e. 0,1,2,3,4,5, and variation in numbers of transmission is typically large. Importantly, there is a small, but finite, probability of leaving zero infections next generation; in the case “genetic extinction” has occurred i.e. the allele has been lost from the populations. The risk of genetic extinction is greatest when there are small number of one type of allele. This is most notable for new mutations, which are by definition, present at a single copy when they first arise; in that case, even with a selective advantage of 5% then the mutation will be lost by chance between 90% and 99% of the time depending on the level of heterogeneity in transmission (see discussion in Box 2 of [14]). Hence it is highly desirable to avoid low numbers of infections with any allelic type in the IBM as they may go genetically extinct purely by chance, even if that allele is advantageous over the longer term.
>
> *Genetic drift* is similar to stochastic change described above but occurs at higher numbers. The risk of random genetic extinction has largely passed when there are large numbers of the advantageous allele, but random fluctuations may obscure the underlying selection. Suppose there are 1,000 resistant infections with selective advantage of 5% then on average there will be 1000*1.05= 1,050 next generation. But the stochastic variation described above will still occur and cause chance variation around this expected number. This is termed genetic drift and introduces ‘noise’ that causes variation in the spread of the advantageous allele. A critical point is that at low selection coefficients the drift can completely obscure the dynamics of spread: in effect the ‘signal’ (selection pressure favouring the alleles) is lost in the ‘noise’ (genetic drift). Intuitively, and correctly, the effects of drift become more pronounced at small population sizes i.e. the smaller the number of infections tracked in the IBM, the larger the impact of drift. This is a well-knows phenomenon in population genetics which, for our purposes, sets a limit on the sensitivity of the IBMs to track selection. i.e. for any given size of IBM there comes a point when selection coefficient become so small that it is obscured by drift and the IBM can no longer effectively track the genetics. The impact of this effect is discussed in detail later in section 2.3

A final key genetic concept is that of effective population size, *Ne*, and the “census” population size, *N*. The latter is simply the number of all malaria clones in the population (i.e. of all allelic types) which we will take as the number of infected humans present in the IBMs (ignoring, for convenience, the fact that some infections may be superinfections i.e. consist of several clones). Population genetic theory used to investigate finite population sizes has been developed for idealised, paradigm population that are assumed to have a constant census size *N* and whose members are assumed to have equal reproductive potential. Most real populations, including most infectious disease species, do not fit this paradigm, which led to the concept of an effective population size which is the size of the paradigm population that would have equivalent properties of that of the census size *N* (see [15] for an introduction to *N*, *Ne* and its impact on genetic drift). Importantly, *Ne* in natural populations is invariably much smaller than *N*. Notably there is considerable heterogeneity in the reproductive success of many infections depending on host factors such as the level of host immunity, the number of secondary contacts made by that person (if the disease is directly transmitted) or how often that person is bitten (if the infection is indirectly transmitted by disease vectors such as ticks or mosquitoes). *Ne* also falls as pathogen populations “bottleneck” due to seasonal patterns of transmission (a common phenomenon associated with mosquito-transmitted disease) and large-scale control programmes (such as bed net distribution programs to control malaria) may cause significant reductions in pathogen census population sizes. These factors will substantially reduce *Ne* with important consequences for genetic drift which, as we show later, limits the sensitivity of IBMs to track selection of advantageous alleles with low selection coefficients. The important point is that the number of infections being tracked, *N*, may appear high, but *Ne* may be considerably lower which has a large impact on the IBMs’ ability to quantify the spread of advantageous mutations

The purpose of this manuscript is to describe how these effects of finite population sizes need to be clearly recognised when incorporating genetic selection into IBMs and describe and discuss how their impact may be mitigated.

### 2. Methods and results

We use the individual-based malaria simulation package OpenMalaria to simulate the spread of drug resistance. This is a highly sophisticated IBM of malaria transmission that incorporates factors such as the acquisition of human immunity against malaria infection, local treatment practices, the level of mosquito transmission, and has been widely used to investigate many aspects of malaria transmission and control. The consequence of this sophistication is that it is highly computationally intensive, so makes an ideal test platform to develop computationally efficient methods of incorporating genetic selection. Technical details of how we ran OpenMalaria are given in Supplementary Information but, in summary, we run the simulation over a 10 year burn-in period before introducing the advantageous allele. OpenMalaria outputs the number of inoculations of each allelic type every 5 days as a cumulative total over that period. We extract the proportion of inoculations carrying the advantageous allele from each 5-day time point to monitor the spread of the advantageous allele and it is these data that enter the regression to estimate the selection coefficient. The number of humans to be tracked in OM is user-specified (we track 10,000 otherwise indicated) and we can vary the user-defined entomological inoculation rate (EIR; the mean number of infective bites per human per year) to vary the prevalence of malaria infection; by default we simulate a prevalence of 15% averaged over all ages based on diagnosis by microscopy.

In these simulations we assume the “advantageous allele” is one encoding drug resistance; the same principle applies to all advantageous alleles, but alleles encoding drug resistance allow us to easily alter their selective advantage simply by altering their level of resistance to the drug being deployed. In the following examples, the drug is assumed to be dihydroartemisinin+piperaquine (DHA+PPQ) which is a widely used front-line antimalarial drug. We simulate different selection intensities driving the advantageous allele by varying the resistance level (IC50) to PPQ; increasing the IC50 value encoded by the advantageous allele increases the level of drug resistance and hence defines more highly advantageous mutations (Table 1). This strategy allows us to investigate the range of selection coefficients that covers “typical” values for drug resistance selection which has been estimated at ~0.02 to ~0.12 (see later discussion). Results obtained for IC50 shifts of 1.1× and 1.4× IC_50_ fold are used as illustrative examples (except in part 2.3) as they represent selection coefficients of ~0.02 and ~0.06 respectively using our default prevalence of 15%. Values higher than this gave largely predictable, deterministic results and IC_50_ shifts in between these values of 1.1× and 1.4× were found to behave in an intermediate fashion that therefore added little to the overall picture.

**Table 1.** How increasing the drug resistance level of the advantageous allele increases its selection coefficient, s, and how the magnitude of s may vary according to infection epidemiology. Drug resistance level is defined as the fold increase in IC50 to the antimalarial drug piperaquine compared to the wildtype allele. Malaria treatment and epidemiology is as described as in the main text and the simulations track 100,000 humans. The left column reports selection coefficients obtained when malaria prevalence was 15%, while the right column reports selection coefficients obtained when prevalence was 1.5%; all estimates started from an advantageous allele frequency of 10%.

| Drug resistance level | Selection coefficient (+/- s.e.).<br>15% prevalence | Selection coefficient (+/- s.e.).<br>1.5% prevalence |
| --- | --- | --- |
| **x1.0125** | 0.002 ± 0.0005 | 0.0001 ± 0.002 |
| **x1.025** | 0.004 ± 0.0004 | -0.0004 ± 0.002 |
| **x1.05** | 0.008 ± 0.0004 | 0.002 ± 0.002 |
| **x1.1** | 0.016 ± 0.0004 | 0.004 ± 0.002 |
| **x1.2** | 0.030 ± 0.0004 | 0.004 ± 0.002 |
| **x1.4** | 0.056 ± 0.0004 | 0.005 ± 0.002 |
| **x2.4** | 0.13 ± 0.0004 | 0.017 ± 0.001 |

### (2.1) Methods used to measure and report the dynamics of spread

Figure 1A shows the spread of an advantageous allele in a haploid organism. Most selection processes start from low frequency so most of the timescale taken to reach operationally significant frequencies occurs at low frequencies. Tracking the entire selection period is both computationally intensive and will also suffer from stochastic fluctuations in low allele number as described above. Fortunately, bacteria and malaria are haploid (we know of no selection that occurs in malaria’s brief diploid phase in mosquito oocysts) so dominance between the two alleles is not an issue and the spread can be linearised as shown on Figure 1B. This offers an methodological alternative to measuring spread over long duration at low frequency: it suggests that, providing selection pressure does not alter as a consequence of allele frequency (an important assumption, discussed later) that spread can be measured over a short period of time (computationally convenient), quantified at relatively high frequencies (minimising the impact of stochastic changes in allele frequency), and extrapolated to the whole selection process.

**Figure 1.**
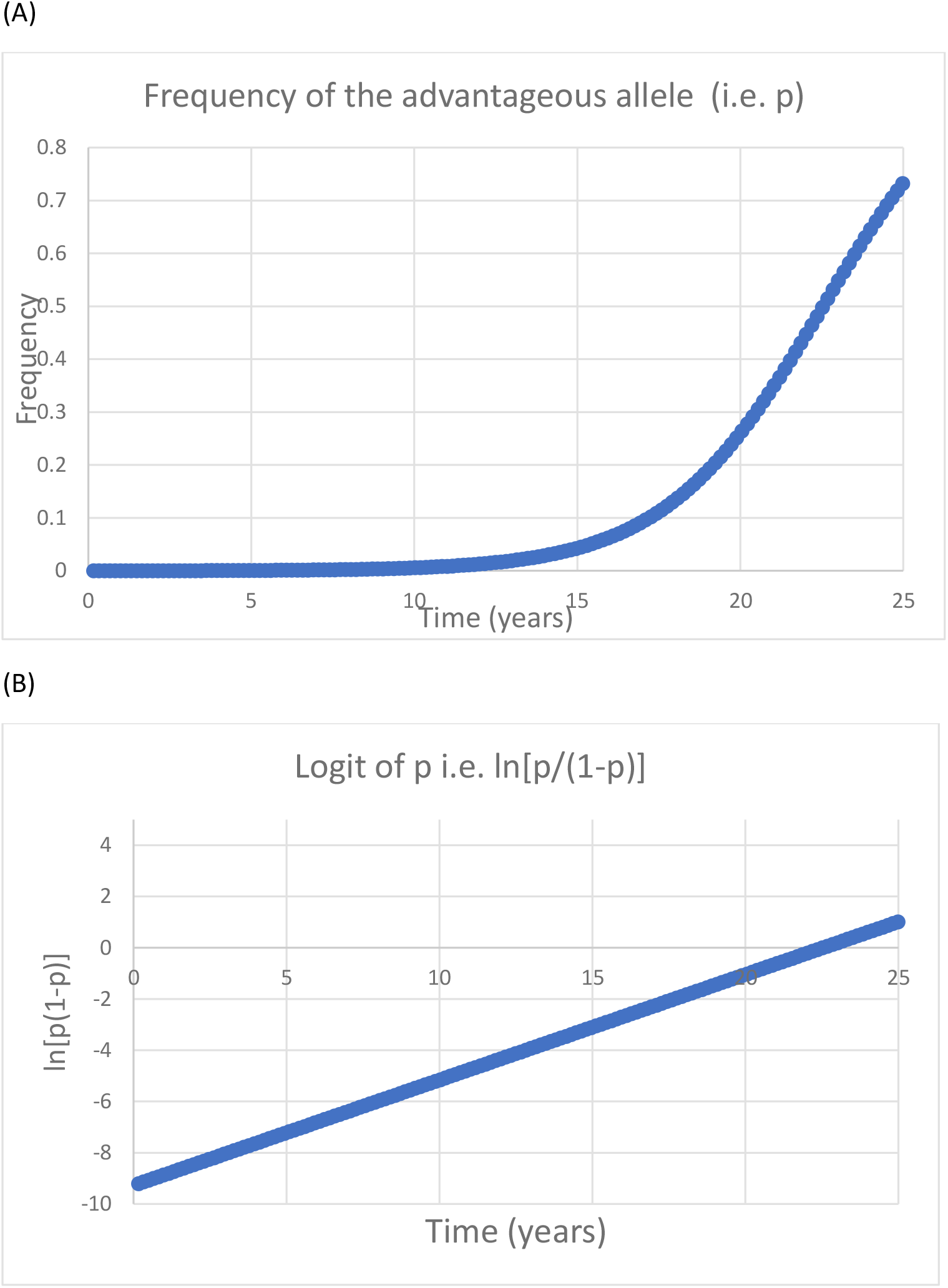
The spread of an advantageous allele in haploid organisms such as bacteria [36] or malaria [17]; this example illustrates dynamics for a selective advantage of s=0.07. Panel (A) shows spread on an arithmetic scale starting from a frequency of 0.0001. Much of the timescale occurs at frequencies that are essentially undetectable (and may be called the period of complacency) before becoming detectable in surveys of a reasonable size. Panel (B) show the same data plotted as its logit, ln[p/(1−p)], where p is the frequency of the advantageous allele and (1-p) is the frequency of the wildtype allele.

We argue that this extrapolation is best achieved by estimating the selection coefficient, usually denoted ‘*s*’, to quantify the rate of spread. This parameter describes the relative ‘fitness’ of the mutant as 1+s compared to the fitness of the wildtype which has fitness 1.0 and is conventionally reported in units of a single malaria generation (see below). Hence, at low frequencies, the mutant spreads at a rate *s* per generation (so if *s*=0.05, then it increases by 5% per generation). Selection coefficient can be easily measured independently of allele frequency, as the slope of ln[p/(1−p)] over time where *p* is the frequency of the advantageous allele (Figure 1B). Computer simulation generally run in “real” time, usually one-day timesteps, so we need to convert the slope of ln[p/(1−p)] from real time to generations. In the current example, this requires an estimate of the duration of a malaria generation. In previous work we have assumed 5 generations per year [16] although other authors used different values, for example Anderson and Roper [17] assumed 6 generations per year. Here we will assume 6 generations per year meaning that each generation lasts 365/6≈60 days so the slope of ln[p/(1−p)] estimated on a timescale of days need to be multiplied by 60 to obtain the selection coefficient. Finally, note that *s* can be negative if the allele is actively being removed from the population (for example by natural selection if the allele is no longer being selected and has a fitness cost) or if the allele frequency decreases due to chance fluctuations (genetic drift); the principles described above apply equally to this situation i.e. its magnitude is measured as ln[p/(1−p)] over time but it will have a negative value (see, for example, Figure 3 of Anderson and Roper [17]).

The dynamics shown on Figure 1 are those predicted by elementary population genetic theory i.e. constant selection pressure in an idealised population of infinite size. Individual-based simulations do not fit this paradigm as selection coefficients may vary over time, most plausibly as a function of allele frequency (see later discussion), and IBMs, by definition, do not track infinitely large populations. All IBMs that we are aware of require a “burn-in” period to allow malaria epidemiology to stabilise in the presence of the human “intervention” (e.g. drug-, RDT-, or vaccine-deployment) prior to introducing the advantageous allele and tracking its spread. This “burn in” period forces researchers to make three key decisions when estimating selection coefficients after introduction of the advantageous allele i.e. what should be the starting frequency of the advantageous allele when introduced into the IBM, when should measurement of ln[p(1−p] start, and how long should measurement last? We illustrate the trade-offs inherent in making these decisions.

Each OpenMalaria simulation was run 100 times for each resistance level with different random number seeds to check consistency of estimates over ‘identical’ runs. Selection coefficients were obtained by linear regression of ln[p/(1−p)] as described above. We set frequency boundaries to avoid the regression tracking small numbers of alleles which would occur at high or low frequencies. The reasoning is that outside these boundaries there may be a relatively small number of alleles of one type and stochastic variation in their number may obscure the deterministic change in their frequency; i.e. the effect of genetic drift as described above. The upper boundary was an advantageous allele frequency (AAF) >90% in all simulations. The lower boundary depended on initial AAF: it was AAF<30% if initial AAF frequency was 50%, AAF<1% if initial AAF was 10% and AAF<0.1% If initial AAF was 5% or 1%; note that the latter two initial AAF values were only every used to produce the data shown on Figure 2. The regression was terminated if the advantageous allele frequency fell outside these boundaries and regression only used data between the initial AAF frequency and the point at which AAF first fell outside the boundaries.

**Figure 2.**
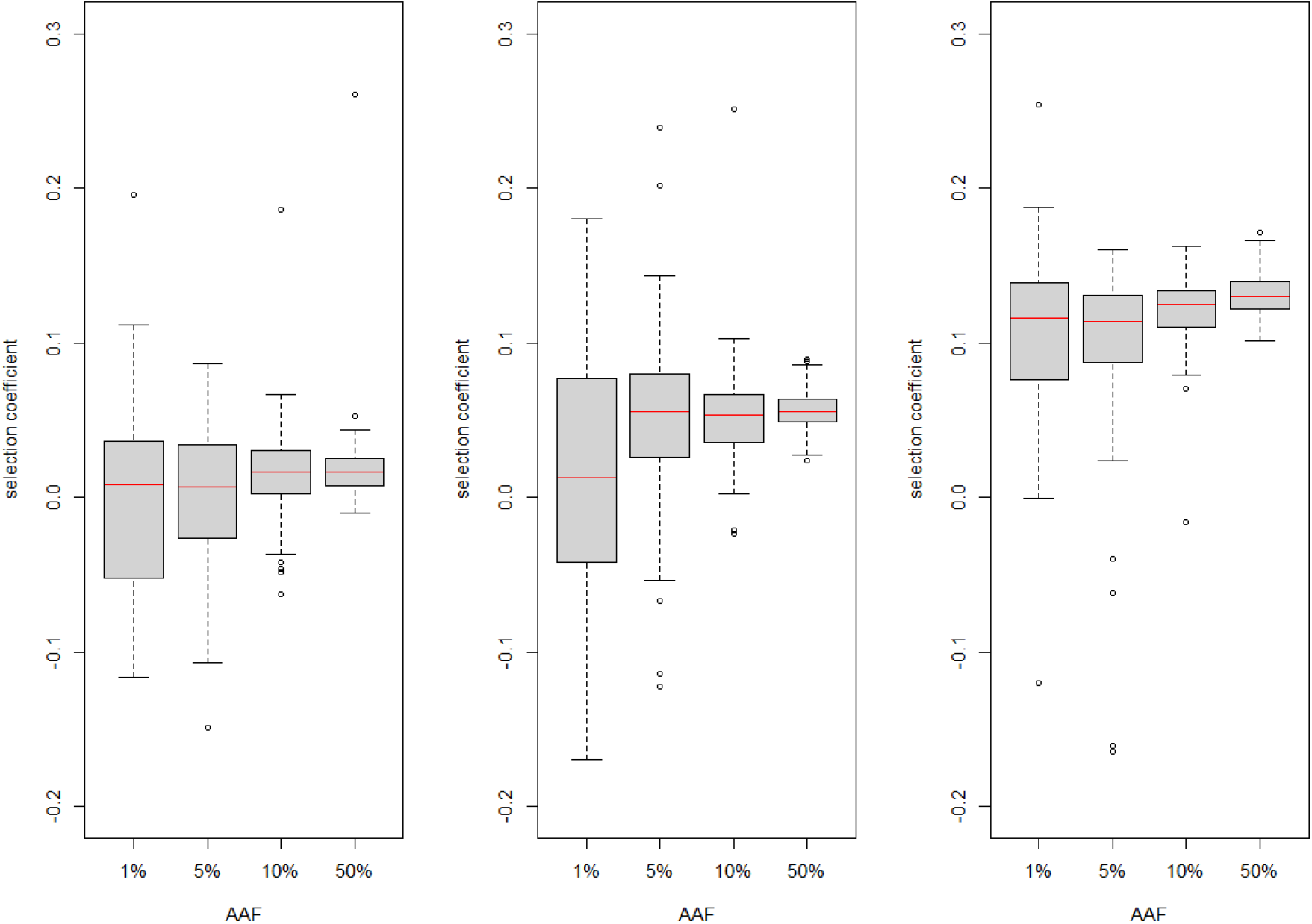
How starting advantageous allele frequency (AAF) affects estimation of selection coefficient, *s*. The boxplots each summarise 96 to 100 estimates except for AFF of 1% where there were 49 estimates (all runs were based on 100 simulations but the advantageous allele fell outside boundaries before regression could be performed for some runs; see SI for more details). Selection coefficients were obtained by simulating 10,000 humans with malaria prevalence of 15% using a regression window that starts 60 days after introduction of the advantageous allele and lasts the next 720 days (assuming AAF stay within bounds, details in main text). Panel (A): the advantageous allele encodes a 1.1-fold increase in drug resistance (s≈0.02). Panel (B): the advantageous allele encodes a 1.4-fold increase in drug resistance (s≈0.06). Panel(C): the advantageous allele encodes a 2.4-fold increase in drug resistance (s≈0.13).

**Figure 3.**
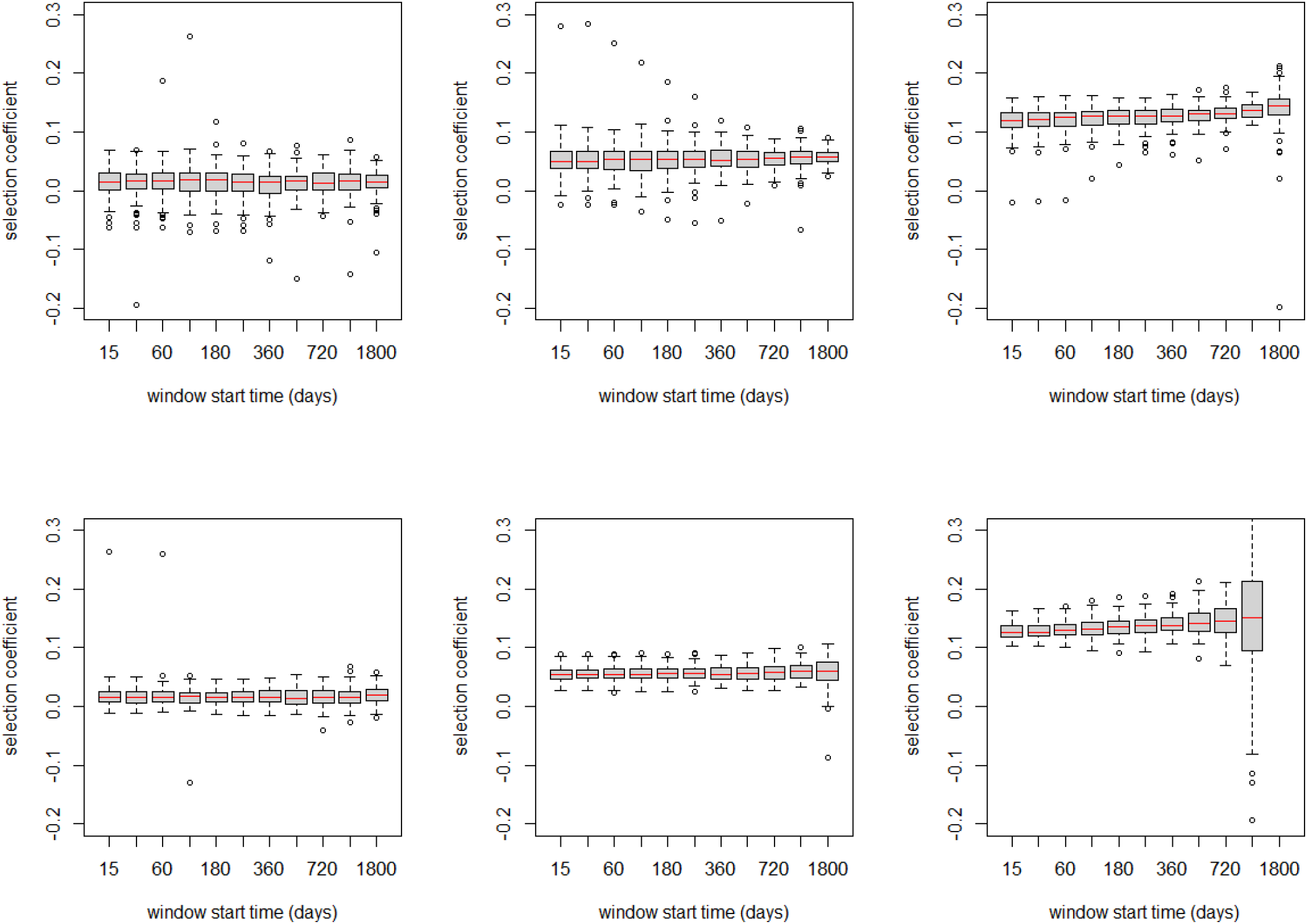
How choice of day (i.e. after introduction of the advantageous allele) used to start the regression window affects estimation of selection coefficients. The X axis shows time post-introduction that the regression window started for values of 15, 30, 60, 120, 180, 240, 360, 540, 720, 1080, 1800 days. The boxplots each summarise 98 to 100 estimates of selection coefficient except where state otherwise*. Estimates were obtained by simulating 10,000 humans with malaria prevalence of 15% using a regression window with duration of 720 days. Left column: the advantageous allele encodes a drug resistance increase of 1.1 fold. (s≈0.02). Central column: the advantageous allele encodes a drug resistance increase of 1.4 fold (s≈0.06). Right column: the advantageous allele encodes a drug resistance increase of 2.4 fold (s≈0.13). Top row is starting advantageous allele frequency (AAF) of 10% and the lower row is starting AAF of 50%. Bottom right panel has two anomalous results: the penultimate column has larger variation because at AAF of 50% and s≈0.13 then our boundary condition of AAF>90% resistance is often exceeded during the 720-day regression window so regression is performed over much shorter periods; the last column is empty because all runs had exceeded the AAF>90% boundary at the time the window started so no regressions could be performed. *Under certain conditions, the boundary conditions were reached before regression could start which reduced the number of runs available for analysis. When starting from 10% AFF with a start time of 1080 the 2.4-fold estimate was based on 84 replicates. When starting from 50% AFF with a start time of 1080 the 2.4-fold estimate was based on 50 replicates. When starting from 50% AFF with a start time of 1800 the 1.4-fold estimate was based on 92 replicates and there were no results for the 2.4-fold increase.

#### (i) Decision #1: What is the appropriate starting allele frequency in the simulation?

Most advantageous alleles start at very low frequencies and most of the timescale of spread occurs at very low frequencies before its clinical impact becomes apparent (the period of complacency [18]) as shown on Figure 1A. Ideally, we would investigate spread at low frequencies, but it is generally extremely difficult to track these very low frequencies in individual-based simulations because of the effect of stochastic variation described above. A biologically reasonable starting frequency might be 10^−5^ meaning 1 in 100,000 infections have the advantageous allele but this is obviously impossible if the simulation, for example, has only 10,000 infected individuals. Even if the simulation did track 100,000 infected individuals, it would still be problematic to introduce an allele at a frequency of 1 in 100,000 as the single infection would exhibit considerable stochastic variation in its subsequent number of secondary, tertiary etc. infections for reasons described earlier i.e., for example, if *s*=0.1 then the infection cannot leave exactly 1.1 offspring but 1.1 will be an average of a distribution with 0,1,2,3,4… secondary infections. The key computational requirement is that small numbers of resistant infections need to be avoided. So a decision needs to be made on starting frequency that ensures a reasonable number of each type of allele such that stochastic variations due to genetic drift are minimised (i.e. the dynamics are largely “deterministic” in the population genetic jargon).

The dynamics of Figure 1 assume constant selective advantage over time. This assumption may hold at very low frequencies where small changes in frequency, say from 10^−6^ 10^−5^, do not significant affect malaria epidemiology. Once frequencies become larger, the epidemiology may begin to change because of the presence of the advantageous allele; for example. spread of a *hrp2*-deletion allele might allow malaria to resurge, treatment rates to fall, and prevalence to increase. We show how selection coefficients may differ depending on epidemiology in Table 1 which shows the impact of a large, but illustrative, 10-fold reduction in malaria prevalence. The selection coefficient is almost 10 times higher when malaria prevalence is 15% than when prevalence is 1.5%. The most likely explanation is that the percentage of people with detectable drug level is 35% in the 15% prevalence group falling to 8% in the 1.5% prevalence group. The proportion of people in the population with sub-therapeutic drug levels is an important driver of resistance (see Box 1 of [19] and Box 2 of [20] for discussions) so higher selection coefficients are generated when there are higher proportion of patients with detectable drug levels. A ten-fold change in prevalence is obviously a very large epidemiological change but serves to illustrate our point about the potential impact of changing epidemiology on estimates of selection coefficient.

There are therefore two conflicting considerations in choice of initial advantageous allele frequency. Ideally, it should be as low as possible to reflect the condition under which most selection occurs (Figure 1) but this is weighed against the benefits of higher frequencies in reducing stochastic noise in allele spread. We examined different values of starting frequencies as shown in Figure 2 and values of 10% and 50% are also compared in Figures 3, 4 and SI Figure S3. We decided on 50% as the default value to minimise stochastic change because Figure 2 shows it greatly reduces the variation in estimating selection coefficient. Note, however, that when values of s are sufficiently high that they reliably dominate genetic drift (s>~0.1 in our default simulations based on 10,000 humans with 15% malaria prevalence) then an initial frequency of 10% can be used; this allows regression to be measured before advantageous allele frequency exceeds the 90% threshold applied in our analyses. We discuss the implications of this choice of initial frequency later and recommend that researchers explicitly discuss how their results obtained at high frequencies are likely to also apply to selection at much lower frequencies.

**Figure 4.**
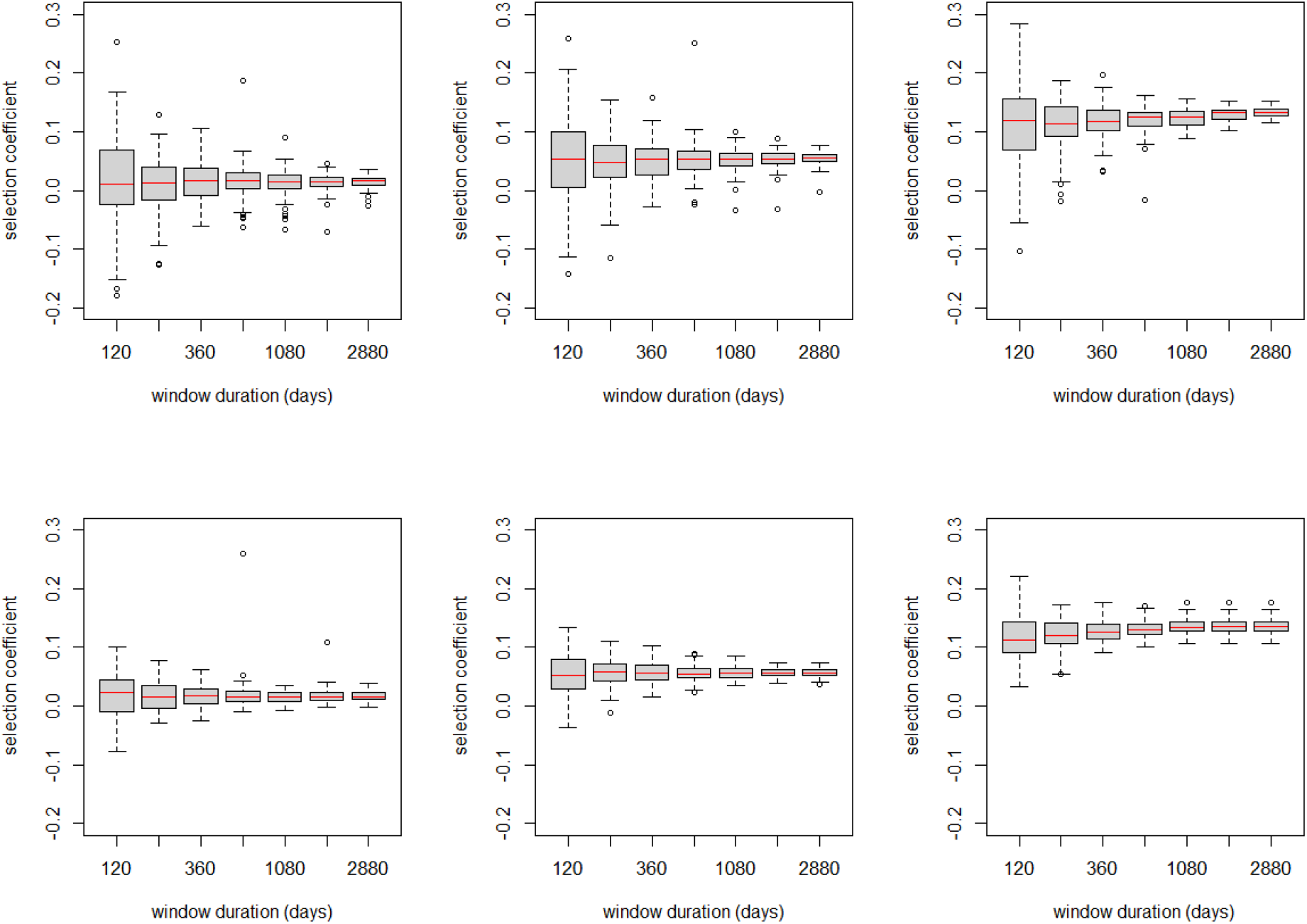
How duration of regression window affects estimation of selection coefficients. The X axis gives duration of regression window and values of 120, 240, 360, 720, 1080, 1880 and 2880 days were investigated. The boxplots each summarise 98 to 100 estimates of selection coefficient obtained from a simulation of 10,000 humans with malaria prevalence of 15% using a regression window that starts 60 days after introduction of the advantageous allele; Left column: the advantageous allele encodes a drug resistance increase of 1.1 fold (s≈0.018). Central column: the advantageous allele encodes a drug resistance increase of 1.4 fold (s≈0.058). Right column: the advantageous allele encodes a drug resistance increase of 2.4 fold (s≈0.13). Top row is starting advantageous allele frequency (AAF) of 10% and the lower row is starting AAF of 50%.

#### (ii) Decision #2: How long after introduction of the advantageous allele should we start measuring the slope of ln[p/(1−p)]?

Decision #1 means we have to introduce the advantageous allele at a relatively high frequency which will inevitably change disease epidemiology, so the second decision is how long after the introduction of the advantageous allele should we start to measure ‘*s*’? The best policy would be to start measuring selection as soon as possible after its introduction to minimise the impact of any epidemiological changes. Plausible start times could be immediately after introduction, after 20 days (to allow secondary infections to be acquired by the mosquito, develop in the midgut, be inoculated and become patent in humans), 60 days (the estimated malaria generation, see above), and so on. We investigated this by varying the start time and quantifying its effect on estimated selection coefficients. Example results using the default duration of ln[p/(1−p)] regression of 720 days are shown on Figure 3 but the results were fairly consistent across other durations of regression (Figure S1). Start times >60 days do not appear to bias the median estimate of ‘s’ nor does it make a big difference to the coefficient of variation around the estimates. We therefore selected a delay of 60 days, equivalent to a single malaria generation, to allow transmission to stabilise.

#### (iii) Decision #3: How long should we regress the slope of ln[p/(1−p)]

The aim is to make the duration of regression sufficiently short that the epidemiology over that time period has not changed significantly due to the introduction of the advantageous allele, while ensuring that the duration is sufficiently long that accurate estimates of ‘s’ can be obtained from the regression. We investigated this effect by altering the duration of regression and quantifying how it affects the estimation of selection coefficients, see Figures 4 and lower panel of Figure S2, assuming, as justified above, that the regression starts 60 days after introduction of the advantageous allele. The duration does not appear to bias the mean or median estimate of ‘s’ but did affect the variation in the estimates, the variation decreasing as duration increased (Figure S2), presumably because more datapoints enter the regression. We chose 720 days as the duration of our regression as it returns stable estimates of ‘s’ with relatively small coefficients of variation (Figure 4).

Finally, We needed to confirm that spread of the advantageous allele had no significant impact on epidemiology (or, if it did, that it did not alter the magnitude of ‘ s’) so that the value of ‘s’ from our regression will be a valid approximation for that occurring during the critical period of spread from low initial frequencies. Figure S1 shows that the estimated selection coefficient did not systematically change with time since introduction of the advantageous allele; frequency will have increased over this time implying that estimates of selection coefficient were not affected by allele frequency.

#### (iv) The impact of these decisions

There appear to be no objectively “correct” solutions to these three decisions, all of which incur trade-offs. It may be possible to tailor the decisions to specific circumstances. For example, if selection coefficients are known to be large, and it is computationally feasible for the simulation to track a large number of infections in a reasonable time frame, then the advantageous allele may be introduced at a much lower starting frequency to minimise its epidemiological impact after its introduction. We wished to avoid using different methods in different circumstances and wanted to identify a robust method which we believe will be applicable over the wide ranges of parameter spaces explored by OpenMalaria, hence we settled on the above methodology. We are not prescriptive on the methodologies to be employed but herein describe what trade-offs we encountered and how we addressed them; importantly, these will almost inevitably occur when tracking genetic selection in IBMs of most infectious diseases. We used in OpenMalaria as a case study in decision making which people using other IBM for other alleles, may find useful.

In summary, the default method we use for estimating ‘s’ in OpenMalaria is as follows. The advantageous allele is introduced at 50% frequency (with the caveat that if ‘s’ is very high, starting frequency of 10% gives a longer duration of regression before frequency exceeds our limits of 90% which may improve accuracy). The regression then starts 60 days after the introduction of the advantageous allele (this one-generation time lag allows malaria transmission stages, gametocytes, to mature and reflect the preferential transmission of the advantageous allele) and continues over 720 days (2 years) which, assuming 6 generations per year (see below) is 12 malaria generations. OpenMalaria outputs data every 5 days, so the full regression period of 720 days provides 144 data points for the regression. We then run repeated simulations and check the boxplot of our estimates (figure 2 to 4) or means and coefficients of variation (Figure S1 to S3) to confirm that we have reasonably stable estimates of ‘s’.

#### (2.2) Incorporating mutations into the IBM

We argued that selection coefficients be measured when advantageous alleles are present at high frequency (to minimize the impact of genetic drift) and used to extrapolate spread of the advantageous allele over the whole period of selection. This period of selection can start from any frequency, so a key question is what starting frequency is most appropriate for this extrapolation. In some circumstances we may be able to simply assume that advantageous alleles are already present at a given initial frequency, for example at a mutation/selection equilibrium. In these circumstances, it is straightforward to use the estimate of ‘s’ to track their subsequent spread from their initial, presumably very low, frequency. For example, if the initial advantageous allele frequency is p(0) then the frequency after ‘t’ generations, p(t) can be obtained by substitution as in the following equations. Figure 1B shows that

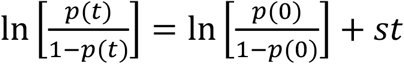

Or, equivalently,

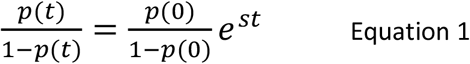

Noting that if the odds of p(t) are ‘x’, i.e. p(t)/[(1−p(t)]=x, then the allele frequency at ‘t’ is given by p(t)=x/(1+x)].

Or, alternatively, the time taken to reach a frequency of p(t) from p(0) can be obtained by substitution as

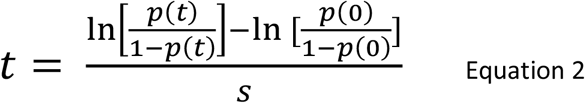

Similarly, the selection coefficient can be obtained from field data reporting frequencies at different times (a common form of data; see later) as

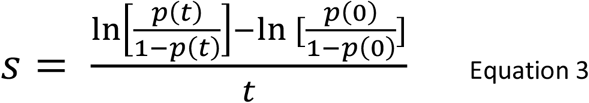

Alternately, the assumption may be that the advantageous alleles are not yet present in the malaria population, in which case their *de novo* input by mutation must be incorporated into the IBM. The problem then is to track the origin and spread of such mutations while avoiding having very low numbers of the advantageous allele within the IBM. We suggest the following strategy to allow the input of *de novo* mutations into the IBM as a 4-stage process.

(i) Define “mutation rate”, μ, as a user-defined input into the simulation We define this mutation rate per inoculum, μ, which in our malaria IBM is the probability that a single mosquito bite delivers an inoculum consisting solely of parasites containing the advantageous allele. This allows details of how mutations arise to be studied external to the IBM as discussed later. Importantly, quantifying the input of new mutations as μ simplifies the introduction of mutations into IBMs, and reduces the stochastic element of their input and improves comparability between replicate runs.

(ii) Use the selection coefficient obtained above to calculate the probability that the mutation successfully survives chance extinction in the first few transmissions after its introduction. This is discussed in more detail elsewhere [14] but the basic results is that if reproductive success follows a Poisson distribution then the probability that the *de novo* mutation survives to become established in the population is approximately 2s when s<~0.1 [21–23]. Reproductive success is generally not Poisson and is much more over dispersed; if we quantify this overdispersal by a negative binominal distribution, as is commonly used in infection transmission, then probability of establishment may be reduced substantially (Table 1 of [14] and see Parsons el al [24] for a more sophisticated discussion). We denote the probability of a novel mutation surviving chance extinction as ψ and can estimate it as

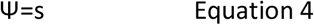

Based on Table 1 of [14] which shows this relationship holds for values of s from ~0.01 to~ 0.1 and a negative binomial distribution of transmissions (a characteristic of most parasite infections) with a dispersal parameter of 0.1 (moderately over-dispersed). Alternative estimates of ψ can be obtained by simulation under different assumptions of dispersal parameter [14], but this simple relationship serves its illustrative role here.

(iii) We can then calculate the expected number of new mutations per (malaria) generation, θ, as

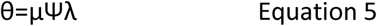

Where λ is the number of successful transmissions in the IBM per mosquito generation. In epidemiological terms this is known as the force of infection (FOI). This is typically much lower than the infective contact rate because many contacts do not produce a viable infection due to factors such as low inoculum size, acquired immunity in the human, etc. In vector-borne disease the contact rate is usually quantified as the entomological inoculation rate (EIR) i.e. the average number of infective bites per human per time period. In the specific case of malaria, Smith and colleagues [25] established a relationship between EIR and FOI which, since malaria EIR is commonly estimated in the field, enables IBMs of malaria to be calibrated against field data when attempting to incorporate mutational input.

Equation 4 enables us to calculate the distribution of “wait times” until a mutation successfully enters the population; rather counter-intuitively to some readers, this is an exponential decay, rather than a normal distribution, see Figure 2 of [14] i.e.

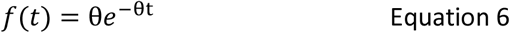

Sampling from this distribution provides the time (in generations) until a mutation first successfully enters the IBM.

(iv) Having obtained a wait time, we can then use ‘s’ to calculate its subsequent spread until it reaches any given point e.g. time to reach 5% frequency. Assume, for example, that the IBM is tracking a population of 1,500 infections (note it is number of infections, not number of humans) then the initial frequency p(0) of a de novo mutant is 1/1,500. The distribution of times for a de novo mutation to successfully enter the population and subsequently reach any given frequency p(t) can be simply obtained by summing the wait time for its successful input into the population (sampled from Equation 6) and the time for its subsequent spread to p(t) (Equation 2). The implicit assumption in this calculation is that the rate of de novo mutations entering the population is small compared to the selective advantage (i.e. θ <<s), so that the spread of the first mutation dominates any later input of new mutations, meaning that we need only track the spread of the first mutation. If this is not the case, the calculation for spread can still be made but allowing recurrent, sporadic de novo mutations to enter the population using Equation 5.

There are two clear advantages of this four-stage strategy of incorporating mutations. Firstly, it allows biological details of how mutations occur to be removed from within the IBM which allows better comparability between studies. Authors may differ in their underlying assumptions about how mutations arise, so this strategy removes such differences from the simulations. Secondly, it removes the temporal variation in wait-times for mutational input from the IBM and places it in Equation 5. It is obviously easier and much faster to repeatedly sample from the distribution in Equation 5 than to repeatedly run replicates of the IBM to capture and quantify this stochastic variation in mutational input.

### (2.3) Identify the power of the IBM to accurately estimate small selection coefficients

In Part 2.1, we deliberately focussed on selection of sufficient strength that selection was able to dominate genetic drift in our simulations i.e. selective coefficients >2% and a malaria population census size, *N*, of ~1,500 (i.e. 10,000 humans tracked with a malaria prevalence of 15%).

IBMs typically simulate humans. If disease prevalence is low as, by definition, occurs in near-elimination scenarios, then the pathogen population size may be small. Suppose a simulation of 10,000 humans with a pathogen prevalence of 1% gives a pathogen population size of 100. This may seem large but, as described earlier, it is effective population size, *Ne*, that is important rather than the observed or “census” population size, *N*. The reason this is so important is that *Ne* sets the sensitivity of the IBMs to detect selection acting on advantageous alleles. As *Ne* becomes small, random frequency changes due to genetic drift become so large that they can obscure selection processes. There are various approximations for deciding whether selection or drift will be the dominant dynamics driving changes in allele frequency but, generally, drift dominates when s<<1/*Ne* and selection dominates when s>>1/*Ne*, and both play important roles when s and 1/Ne are around the same magnitude. Suppose selection coefficient is 0.02 which is moderately strong i.e. a 2% increase per generation, this suggest it will only dominate drift when Ne>>50.

The problem is that, except in very simple cases, there is no algebraic means of converting the “census” population size observed in IBMs, into the equivalent *Ne*. One option would be to track neutral (i.e. non selected) genetic markers in the simulation and use the level of linkage disequilibrium to estimate *Ne* (e.g. [26]). However, it is probably more informative to empirically estimate the limit of sensitivity of the IBM to detect small selection coefficients. We demonstrate this by further reducing the resistance level (IC50) of the advantageous alleles below that investigated in Part 2.1 i.e. fold increases in IC50 are less than 1.1 (Table 1). The dynamics of spread when selection coefficients are low is shown on Figure 5 and shows the coefficient of variation around mean estimates of ‘s’ (cf Figures 3 and 4 where selection is higher and dominates drift). Figure 5 has the following pattern:

The top line was obtained from the following simulation:

- 10,000 patients with malaria prevalence of 15% and starting mutant frequency of 10% *N*=1,500

There are then two lines that are effectively superimposed i.e.

- Increasing the starting advantageous allele frequency to 50% this increase sensitivity compared to the top line, presumably by reducing stochastic variation is allele number.
- Increasing population size to 100,000 and decreasing malaria prevalence to 1.5% this gives the same malaria census population size as 10,000 people with 15% prevalence (i.e*. N*=1,500) so it is an excellent demonstration that it is the malaria population size being tracked *not the human population size*, that is the critical factor governing ability of the IBM to estimate selection coefficients.

Finally, there is a lower line obtained from the following simulation as follows

- 100,000 patients with malaria prevalence of 15% and starting mutant frequency of 50%. This has N=15,000 i.e. a 10-fold greater malaria population size compared to the 100,000 population with 1.5% prevalence. As expected, the ability of the IBM to accurately estimate selection coefficients is greatly increased.

**Figure 5:**
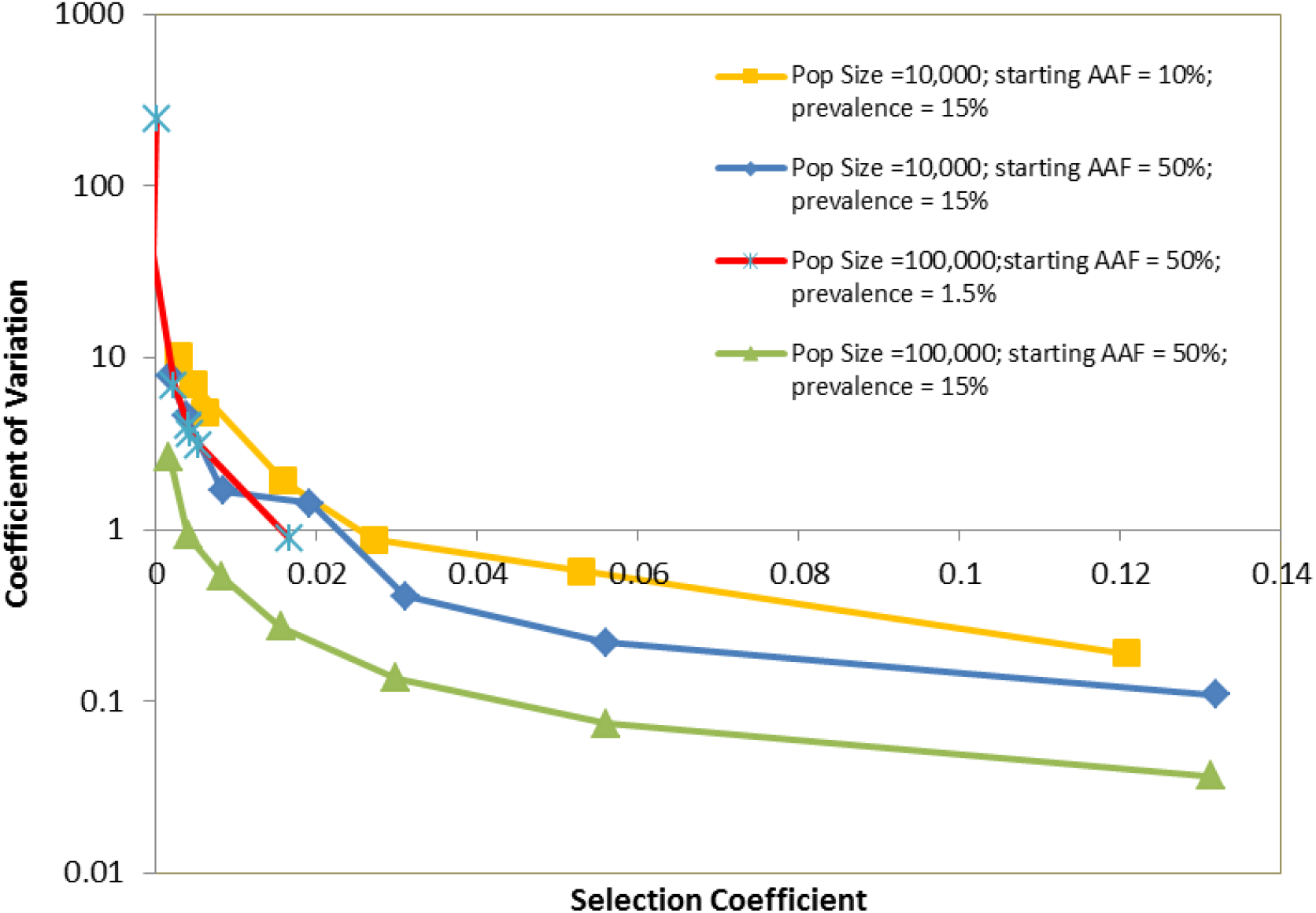
How the power of a simulation to detect small selection coefficients depends on its size. Four parametrisations were investigated

- The default simulation size i.e. 10,000 humans with a malaria prevalence of 15%; starting advantageous allele frequency (AAF) of 10%
- The default simulation size used in this work i.e. 10,000 humans with a malaria prevalence of 15%; starting AAF of 50%
- Simulation of 100,000 humans with a starting AAF of 50% and malaria prevalence of 1.5%, meaning malaria population size is nearly identical (there will be stochastic differences) to the 2^nd^ parameterisation where population size is 10,000 but prevalence is 15%.
- Simulation of 100,000 humans with a starting AAF of 50% and malaria prevalence of 15% i.e. a 10 fold increase in malaria population size compared to the default size of 10,000 Each parameterisation was run 100 times and selection coefficient estimated by regressing over 720 days starting 60 days after introduction of the advantageous allele. Note that the third parameterisation (red line) is associated with lower selection coefficients (see Table 1) and consequently is only plotted at the extreme left of the X axis.

As expected, the accuracy of estimation does increase as the number of infections tracked is increased, either by increasing the number of patients tracked and/or by increasing malaria prevalence (Figure 5). In the specific case of default simulations of 10,000 humans with 15% malaria prevalence (blue line of Figure 5), they are unable to accurately estimate values of ‘s’ less than about 0.02 because the CV increases to the extent that a large number of replicates would be required to get an accurate standard error around the estimated value of *s*. Increasing population size to 100,000 enables estimates of s down to around 0.005 (green line of Figure 5). We base this limit on a CV<~1 but this is arbitrary and illustrative and, in reality, the limit depends on how many replicates can be feasibly run. A larger number of runs will reduce standard error around the estimates but, importantly, it may be more computationally efficient to run fewer, large simulations, rather than a larger number of small simulations. Recall that standard error around the mean (SEM) is given by

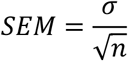

Where σ is the standard deviation of estimates and n is sample size. If we take the central point plotted in Figure 5 (i.e. s=0.0156; see Table 1) with 50% starting frequency and 15% prevalence, then a population size of 10,000 has a CV of 1.43 (figure 5) resulting in σ =0.022 and a population size of 100,000 has a CV of 0.27 (figure 5) resulting in σ =0.0042. Assuming the population simulation scales (i.e. a run tracking 100,000 individual requires 10 times more run-time than a population of 10,000) then a smaller number of simulations tracking a larger population provides better accuracy. For example, 100 simulations of 10,000 individual gives SEM of 0.0022 while 10 runs of 100,000 gives a SEM of 0.0013. Similarly, 50 runs of 10,000 gives a SEM of 0.0032, while 5 runs of 1000,000 gives a SEM of 0.0018. It is therefore likely that running exploratory trial runs to establish the sensitivity of the IBM to detect values of *s* around the anticipated value, could lead to optimised IBM simulation strategies that can significantly reduce the standard errors associated with parameter estimates.

‘The main factor that reduces *Ne* below the census size is the differences in reproductive success of different infections (in malaria, most plausibly due to heterogeneity in mosquito biting rate and acquired human immunity affecting an individual’s infectivity and susceptibility). Adding factors that increase this heterogeneity will further reduce *Ne* and hence the ability to detect low magnitudes of *s*. Consequently, the ability of IBMs to detect low values of *s* is not simply dependent on population size, but on their structure and calibration; this emphasises the need to continually check sensitivity of IBMs before drawing conclusion of magnitude of selection coefficient.

## 3. Discussion

IBMs for complex disease such as malaria have an intrinsic trade-off: the more realistic and complex the biological and clinical description of transmission and disease, the more computational power is required to simulate populations, and the smaller the number of patients/infections that can be tracked in a reasonable timeframe. This is not usually an issue for simulating the overall epidemiology (although stochastic loss of the pathogen population may occur at low prevalences), and IBMs of malaria have been successfully used to investigate a number of control measures as described in the Introduction. However, the finite sizes of IBM do affect their ability to incorporate genetic selection of advantageous alleles. In the specific case of malaria populations, such alleles evolve to counter control interventions such as those encoding drug resistance, vaccine insensitivity and RDT escape-mutations. Given the likely impact of these alleles, a strategy to incorporate such genetic processes into IBMs is urgently required and to our knowledge this is the first attempt to explicitly do so. We make four specific recommendations that may help other researchers attempting to do this

### Recommendation #1. Report selection coefficients and validate the values obtained from IBM against field data

We strongly recommend that IBMs be interrogated to obtain the selection coefficient, s, of the advantageous allele. This is a common scale which can be used to compare results from different IBMs and which can be used to calibrate/validate against field data which typically report selection coefficients acting on alleles (see drug resistance examples below). Selection coefficients also affect other genetic process as was briefly described elsewhere [27] i.e. they determine rate of geographical migration rate, survival probability of mutations (as described above), the genetic impact of selective sweeps and determines the frequency of resistance prior to the introduction of a novel drug as a mutation/selection balance.

As an example of IBM validation using selection coefficients, we searched the literature for estimates of selection coefficients associated with alleles encoding malaria drug resistance in malaria and found these range from around 0.02 to 0.12 as in the following examples (which we do not claim to be exhaustive). Anderson & Roper [17] reported selection coefficients of 0.05 and 0.076 for two *dhfr* alleles and ∼0.13 for a *dhps* allele. Anderson [28] collated data suggesting values of s ranging from 0.03 to 0.11 for the same mutations. Estimates from SE Asia are s~=0.11 for *dhfr* [29] and s~0.08 for *kelch*13 [30]. Nwakanma and colleagues reported values of *s* as 0.15, 0.13, 0.11 and 0.11 for *crt*, *mdr*, *dhfr* and *dhps* respectively over a 25 period in Gambia; note that they assumed 2 generations per year so, for consistency with the assumption of 6 generations per year, these values should be divided by three giving estimated in the range 0.03 to 0.05. Nsanzabana et al [27] reported values of *s*=0.02 to 0.6 for 3 loci undergoing selection over a 12 year period in Papua New Guinea. IBMs producing values in this range are re-assuring while those producing higher or lower values may require supporting clarification about why they produce results different from field observations.

Some authors report basic genetic outputs from the models (e.g. Watson et al [31] report the spread of *hrp*2-deletions from various starting frequencies; their Figure 1) while some do not report the dynamics of mutational spread and simply report epidemiological/clinical impact (e.g. [32]). Simple description of genetic spread and/or clinical impact are informative but we argue strongly that the dynamics should also be described by a selection coefficient as this allows a reader to easily calculate spread from his/her own choice of initial frequency and is a common genetic scale on which to compare selection processes. Selection coefficient can be extracted from previous work using our approach. For example, Watson et al [31] show the increase in frequency of *hrp*2-deletion over time in their Figure 1. The change in frequency between values of 0.25 to 0.75 appears linear for most of their plots (in line with our results, figure 1A of this paper) and generally takes 5 years, equivalent to 30 malaria generations, to spread from 0.25 to 0.75. Substituting these values into our Equation 3 (i.e. p(0)=0.25, p(t)=0.75, t=30) gives a value of s=0.07 which is highly consistent with ‘s’ observed for drug resistance, presumably because the underlying selective forces are similar i.e. drug resistance mutations survives treatment while *hrp2*-deletions simply avoid treatment. This example clearly shows how reporting ‘s’ can effectively summarise a whole graph of data (i.e. Figure 1 of [31]) while also allowing easy comparisons between studies.

We have reported selection coefficient in units per generation, even although OpenMalaria runs in five-day timesteps with overlapping generations. The per generation timescale is a population genetics convention but can be easily interconverted to units per day, or perhaps more convenient, per year (it only has to be in units per generation in this work for application of Equation 4). We do however, strongly recommend that selection coefficients be reported even if their estimation is not the primary objective of the study For example, Nguyen et al [32] reported how selection of malaria drug resistance affected clinical outcomes such as treatment failure rates but did not report underlying selection coefficients; if these could have been extracted and reported it would have helped comparability between studies, and also demonstrated that the dynamics were consistent with field estimates describing selection for resistance.

### Recommendation #2. Use the magnitude of ‘s’ to optimize the computational approach

Any IBM of finite size will inevitably have a lower limit below which ‘s’ cannot be estimated with precision; see Figure 5. A first step would therefore be to empirically estimate its power and avoid trying to obtain estimates of ‘s’ below this lower limit. This is particularly relevant where extensive explorations of parameter space are planned, as it may be inevitable that small values of ‘s’ may be encountered and need to be avoided, or possible discounted in subsequent analysis as being unreliable. In these circumstances it is also beneficial to optimise the computational approach as detailed earlier i.e. decide whether it is more computationally efficient to run a smaller number of replicates of larger IBMs, or vice versa.

### Recommendation #3. Consider how best to incorporate mutations

If mutations are already present in the population before deployment, presumably in mutation/selection balance (e.g. for malaria see Equation 2 of [13]) they can be simply incorporated as starting frequency p(0) in Equations 2 and 3. However, if new de novo mutations are to be incorporated into the IBM, it is best to try and make details of how mutations arise external to the simulation and summarise their input into the IBM as simple mutation rate per inoculum. In particular, use of Equation 5 to incorporate wait times for de novo mutation to enter the population is far more efficient than using repeat runs of the entire IBM to achieve the same purpose.

How mutations give rise to advantageous alleles may be relatively non-contentious in some situations. For example, deletions in *hrp*2 or the mutations that result in altered antigenic profiles in vaccine-escape alleles presumably reflect relative well characterised eukaryote deletion rates and codon mutation rate, respectively. In the example of drug resistance mutations in malaria this is extremely unclear, Hastings [14] identifying four possible sources of i.e. already present before drug deployment in a mutation/selection balance, selected *de novo* from among malaria infection at time of treatment, selected *de novo* from new infections emerging from the liver, spontaneous mutations that occur in untreated humans or in the mosquito stages that are subsequently inoculated as resistant. The relative importance of each is unknown and may even vary between drugs (e.g. resistance to atovaquone can be observed being selected from among malaria infections at time of treatment in around 30% of treatments [33]). Hence it makes sense to externalise assumptions about how resistance mutations arise from the IBMs and use the latter simply to track their subsequent rate of spread and likely clinical impact.

### Recommendation #4. Explicitly discuss how selection coefficient measured in the IBM at a given starting frequency is relevant to spread at low frequencies

There is dilemma at the heart of bringing selection into IBMs: the desire to have low advantageous allele frequencies to reflect the epidemiological setting at which most selection occurs (Figure 1), and the desire to have large numbers of each type of allele to accurately estimate selection coefficients (Figure 2).

The most significant compromise we had to make in our simulations was to assume a high frequency (10% or 50%) of the advantageous allele when introduced into the IBM to ensure a sufficiently large number of each allelic type i.e. advantageous and wildtype. Table 1 shows that ‘s’ may vary depending on the underlying epidemiology so the first check we made was to examine epidemiology outputs from OpenMalaria to confirm that epidemiology was not changing rapidly as the advantageous allele (resistance in our case) spreads. The second check was to consider if selection coefficient may rely on allele frequency. For example spread of drug resistance may affect intrahost dynamics to complicate and even stabilising the dynamics of spread (previously discussed in [34]). The dynamics of *hrp*2-deletions provide an excellent example of how ‘s’ may depend on frequency. Superinfection (i.e. simultaneous infection with two or more malaria clones) is common in areas of moderate to high transmission. *Hrp*2-deletion mutants co-infecting humans with wildtype will be common at low frequencies and will presumably have reduced selective advantage because the co-infecting wildtype, *hrp*2-expressing parasite may present a signal sufficiently strong for diagnosis to occur. However, as deletion frequency increases, so will the proportion of superinfections consisting solely of *hrp*2-deletions and this increases their selective advantage as they will escape diagnosis. In this case we would argue that estimates of selection coefficients obtained at higher frequencies *hrp2*-deletion are only valid if superinfection does not occur i.e. there is only a single malaria clones in each host. A suitable secondary check might be to assume a variation in MOI and work out the probability that a *hrp*2-deletion is detected, and then incorporate this probability of detection into the IBM as a second check.

## Conclusions

It is fortunate that malaria, and most bacterial and viral pathogens, are haploid i.e. contain only a single copy of each gene. If diploids are considered (e.g. the diploid worms responsible for human diseases such as elephantiasis, river blindness etc) or insecticide-resistance in the vectors of diseases such as malaria, sleeping sickness, tick-borne relapsing fever, the genetics becomes much more complicated. The dominance relationship between the wildtype and advantageous allele needs to be considered and this means that the spread of advantageous alleles will not be linear on a logit scale (i.e. Figure 1b)

The results presented here were obtained using one specific IBM, OpenMalaria, but the underlying principles are universal not just to malaria, but to bringing genetic selection into IBMs of other infections (see, for example, [35] for a description of a more general, open-source IBM applicable to several important human diseases). The most implementation-dependent recommendation is likely to be the specifics of how to measure selection coefficient i.e. choice of starting frequency of the advantageous allele, how long after introduction of the advantageous allele to start regression, and the optimal duration of the regression. This may vary both with the IBM’s underlying structure and assumptions, and also with its calibrations e.g. how rapidly transmission and epidemiology stabilise after introduction of the advantageous allele. We suggest researchers using IBMs to track genetic spread follow a similar suite of analysis to ourselves. Our recommendation that measurement be delayed for one malaria generation (60 days) and to continue over the subsequent 2 years (i.e. for 20 to 24 malaria generations) appears robust in our simulations and seems intuitive sensible. The second approach, that of incorporating mutation externally to the simulations, should be widely transferrable across IBMs. The final methodology, to validate the IBM’s ability to accurate estimate the small selection coefficient by examining CV over replicates, is also likely to be universal across platforms; it will be highly informative to see how different platforms perform in this respect. What is clear is that it is highly advisable to carry out these types of checks when incorporating genetic selection into IBMs and to report them in the subsequent publications. In particular, the recommendations we list above will apply to all IBM simulations that incorporate genetic selection, irrespective of the disease. We do not imply that the same quantitative decisions will apply to all systems (e.g. our decision to track selection for 720 days starting 60 days after introduction of the advantageous allele) but we do urge other researchers to recognise that such decisions will occur in their simulations either implicitly or explicitly, and that they be addressed and discussed. We have tried to provide an illustrative roadmap for making such decisions and look forward to future work in the topic.

## Acknowledgements

This work was funded by the Malaria Modelling Consortium (grant # UWSC9759 to IH).

