## Supplementary Information for "Incorporating genetic selection into individual-based models (IBMs) of malaria and other infectious diseases"

**This supplementary information has three parts**

**SI.1 Technical details of running OpenMalaria used to obtain the results described in the main text.**

**SI.2. How force of infection (FOI) was extracted from entomological inoculation rate.**

**SI.3. Additional plots supporting our choice of regression window**

**SI.1 Technical details of running OpenMalaria (OM) used to obtain the results described in the main text.**

We use OpenMalaria (OM) as an exemplar individual based model (IBM) that has been widely used to investigate malaria epidemiology and control (a full list of publications can be found at https://github.com/SwissTPH/openmalaria/wiki/References). However, we keep its calibration as simple as possible so that the methodology can be duplicated on other IBM simulation platforms if required.

It is an OM technical restriction that both the advantageous and wildtype alleles must be present from the start of the simulation. We therefore need to set an initial frequency of the advantageous allele at the start of the burn-in and hope that this input allele frequency does not alter too much by chance fluctuations during the burn-in. We refer to this initial frequency as "target frequency and use boundary frequencies (described in the main text) to exclude any runs that have drifted too far from their target frequencies.

OM version 40.1 was used to simulate all runs. The scenario was based on unpublished work examining prevalence after deployment of mass drug administration (MDA) in Zambia with the population demography and vector species simulated being identical to this work (T. Smith, pers. comm). Simulations were run for 30 year consisting of two parts.

- An initial 10 year "burn-in" period. Dihydroartemisinic+ Piperaquine (DHA+PPQ) was the only drug deployed for routine use in the health care system and parasites were fully sensitive to DHA+PPQ. Access to treatment was set to 40% and there was no self-treatment by any antimalarial drug outside of official treatment sources. Patients seeking treatment were tested for malaria infections by microscopy (diagnosis in OM is probabilistic with a 50% chance of diagnosis when parasite density reached 20 parasite/uL; specificity of this diagnosis is 100%) and only patients diagnosed as infected were treated. Importation of malaria infections was present at a rate of 5 infections per 1,000 humans per year; these importations had a resistance allele frequency equal to the target frequency so helped stabilise allele frequencies against genetic drift during the burn-in. Parasite killing by both drugs was modelled using a mechanistic PK/PD framework with DHA modelled as 1-compartment dynamics and PPQ modelled as 2-compartment dynamics using dosing regimens and parameters as described previously in [1].
- A twenty year "intervention" phase. Resistance to PPQ was introduced at the end of the 10 year burn-in period by altering the drug sensitivity of the resistant allele (it previously had a sensitive profile to avoided being selected during burn-in). Importation was switched off to avoid it obscuring the selection process. Subsequent spread of resistance was monitored by recording the proportion of inoculations into the human host for each genotype every 5 days.

The use of the term "intervention" is potentially confusing in this context. Its most obvious interpretation is deployment of a control intervention such as a new drug, deployment of RDT, a vaccination programme, etc . In OpenMalaria an "intervention" is simply a term denoting the opportunity to change the parameterisation of the IBM run. Often these are true, external human interventions, but here we simply use the "intervention" to introduce drug resistance alleles, switch off malaria importation, and monitor their subsequent spread. In this case DHA+PPQ was deployed throughout burn-in, and the only change made at 'intervention' was to introduce the resistant alleles. Similarly, if we had been investigating the impact of *hrp2*-deletions, RDTs would have been deployed during the entire burn-in period and the 'intervention' would have been used to introduce *hrp*2-deletions and track their spread.

Other technical notes for running OM are as follows:

- The number of humans is a direct input into the simulation.
- Malaria prevalence is adjusted by altering the magnitude of the annual inoculation rate.
- Seasonality in malaria transmission is absent.
- We do not allow variation in PK or PD parameters. This is to minimise variation between runs and focus on the interplay between selection coefficients, details of the regression windows, and population size.

Selection coefficients were measured as described in the main text using the regression coefficients estimated using the base R package function lm.

Finally, note that the exact parametrisation is not important for the present purposes. The methodology was designed to be as simple and repeatable as possible because its primary purpose was simply to produce a series of advantageous alleles with differing selection coefficients to mimic a general selection process.

**SI.2. How force of infection (FOI) was extracted from entomological inoculation rate.**

The force of infection (FOI) was required in Equation 5of the main text to incorporate the input of new infections. The annual entomological inoculation rate (EIR) is an input parameter of OpenMalaria and here we show how FOI can be extracted from EIR using the method described by Smith et al [2] as described in their equations 2-7

Briefly, force of infections, λ(i,t), is calculated as shown below:

$$\lambda(i,t) = Sp(i,t)*Ea(i,t) \qquad \text{(Equation S2.1)}$$

where: λ(i,t) = force of infection (age adjusted) and Ea is age-adjusted EIR from simulated EIR based on assuming an average unweighted age of 50 years old. Sp(i,t) denotes the survival function probability that the progeny of each inoculation survive to give rise to a patent blood stage infection:

$$Sp(i,t) = S_1(i,t)*S_2(i,t) \qquad \text{(Equation S2.2)}$$

Where $S_1$ captures innate density dependent effects and $S_2$ measures the effects of acquired pre-erythrocytic immunity. S1 is calculated as follows:

$$S_1(i,t) = S_{inf} +((1-S_{inf})/(1+(Ea(i,t)/E^*)) \qquad \text{(Equation S2.3)}$$

Where, $S_{inf}$ = the lower limit of $S_1(i,t)$ when the inoculation rate is large, and E* is the value of Ea(i,t) at which half the reduction in $S_1(i,t)$ is achieved. In the paper Sinf and Estar are fitted from field/clinical data as 0.049 (unitless) and 0.032 inoculations/person-night, respectively. We use this parameterisation is our analysis.

$S_2$ is calculated as follows:

$$S_2(i,t) = S_{imm} + ((1-S_{imm})/(1+(Xp(i,t)/Xp^*)^{\gamma P})) \qquad \text{(Equation S2.4)}$$

where: Simm= maximum survival of inoculum in immune individuals. Xp* is the critical value of cumulative exposure and $\gamma^P$ is a hill coefficient represent the sharpness of change is $S_2$. In Smith et al [2] $S_{imm}$, Xp* and $\gamma^P$ are fitted as 0.12, 523.0 inoculations and 5.1, respectively. We use this parameterisation in our analysis. Equations S2.3 and S2.4 are substituted in Equation S2,2, which is then substituted in Equation S2.1.

**SI.3. Additional plots supporting our choice of regression window.**

Figure S1. How choice of the day (after introduction of the advantageous allele) that regression starts affects estimated selection coefficients; regression lasted for 120 days (top row) or 720 days (bottom row). The left-hand column shows estimates of mean selection coefficient and the right-hand column shows coefficient of variation (CV) among these estimates. Methods: Mean and CV values were obtained from 100 simulations with different random seeds. Target frequency of resistance was set to 50%. Resistance levels corresponded to a 1.1-fold increase in sensitive $IC_{50}$ value (blue crosses) and 1.4-fold increase in sensitive $IC_{50}$ value (red crosses).

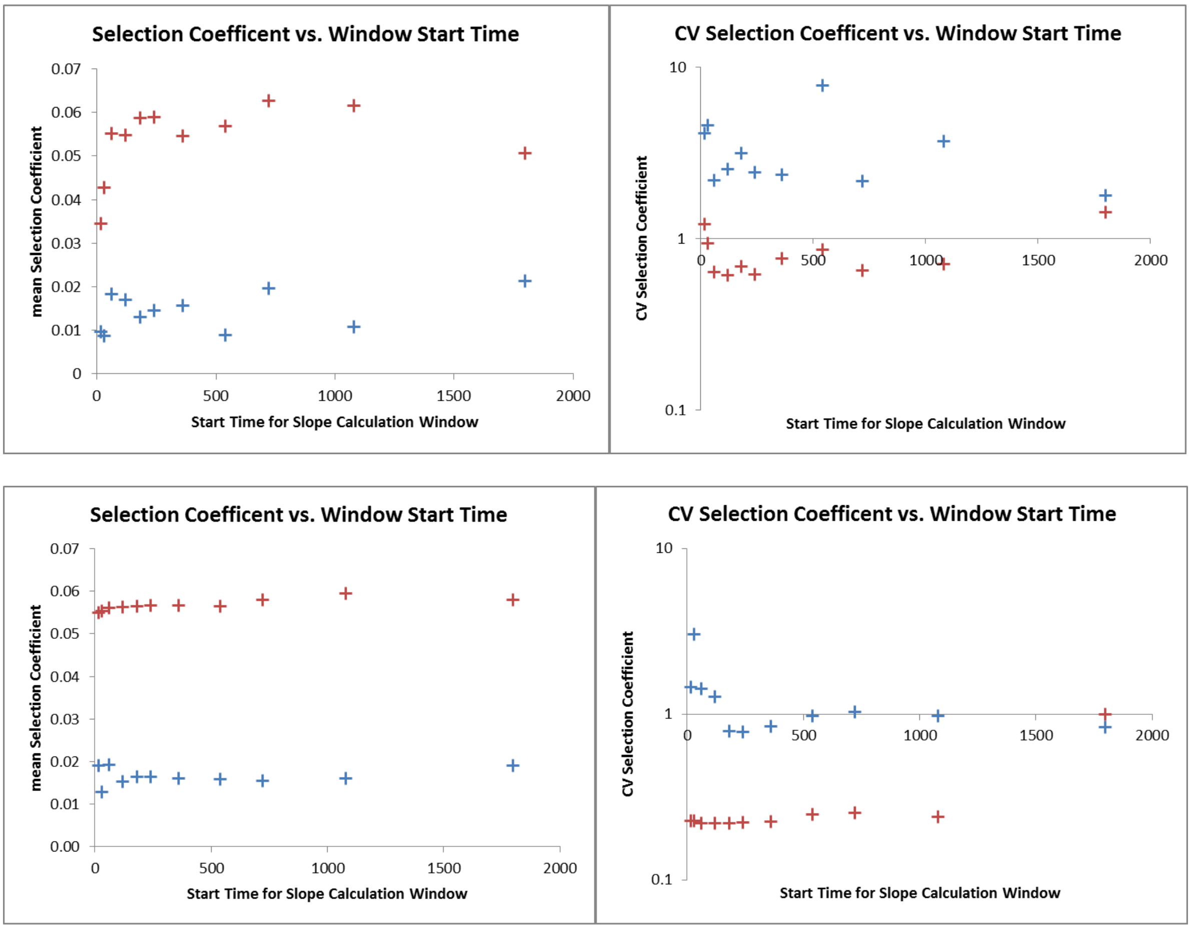

Figure S2. How duration of regression window affects estimated selection coefficients; regression started 15 days after introduction of the advantageous allele (top row) or after 60 days (bottom row). The left-hand column shows estimates of mean selection coefficient and the right-hand column shows coefficient of variation (CV) among individual estimates. Methods: Mean and CV values were obtained from 100 simulations with different random seeds. Target frequency of resistance was set to 50%. Resistance levels corresponded to a 1.1-fold increase in sensitive $IC_{50}$ value (blue crosses) and 1.4-fold increase in sensitive $IC_{50}$ value (red crosses).

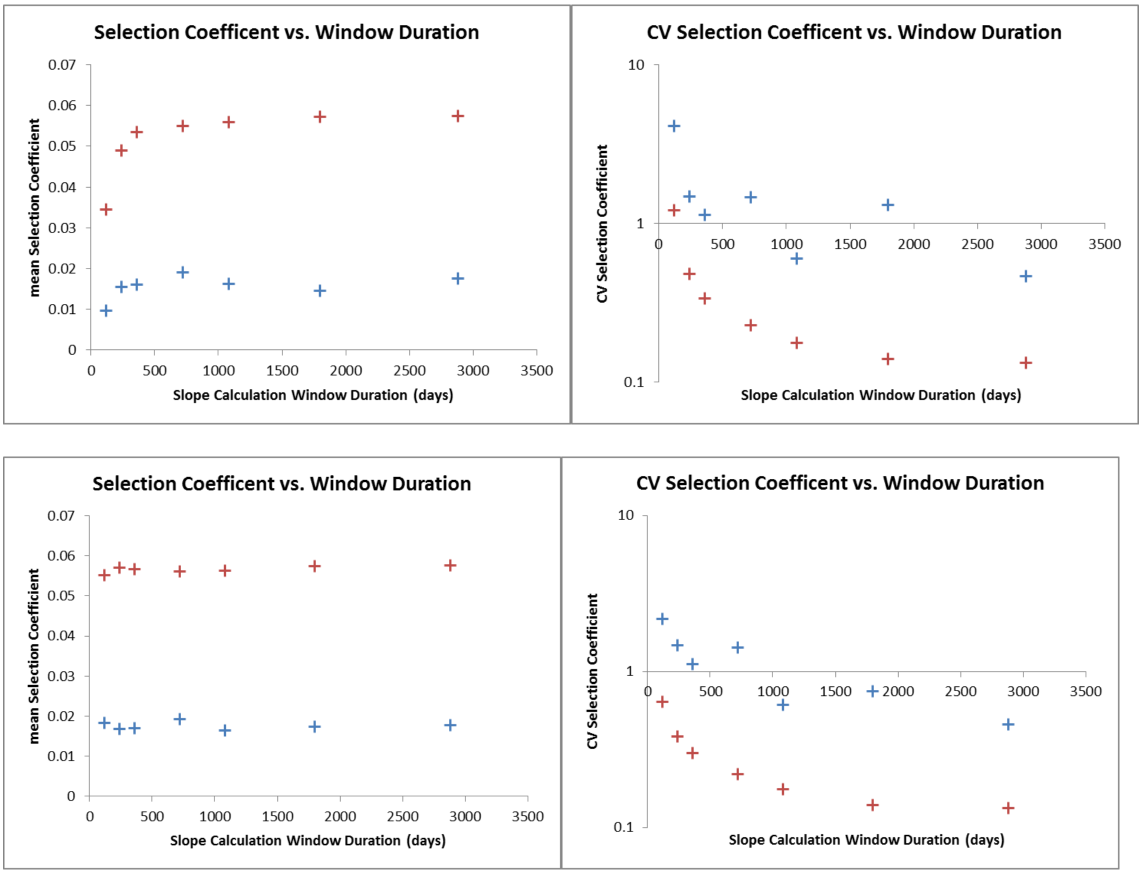

Figure S3. How frequency of resistance affects estimated selection coefficients; target frequency of resistance was set to 10%.

- Top row: as for figure S1, top row, but with 10% (cf 50%) target starting frequency of resistance.
- Second row: as for figure S1, bottom row, but with 10% (cf 50%) target starting frequency of resistance.
- Third row: as for figure S2, top row, but with 10% (cf 50%) target starting frequency of resistance.
- Fourth row: as for figure S2, bottom row, but with 10% (cf 50%) target starting frequency of resistance.

The left-hand column shows estimates of mean selection coefficient and the right-hand column shows coefficient of variation (CV) among individual estimates. Methods: Mean and CV values were obtained from 100 simulations with different random seeds. Resistance levels corresponded to a 1.1-fold increase in sensitive $IC_{50}$ value (blue crosses) and 1.4-fold increase in sensitive $IC_{50}$ value (red crosses).

Fig S3.

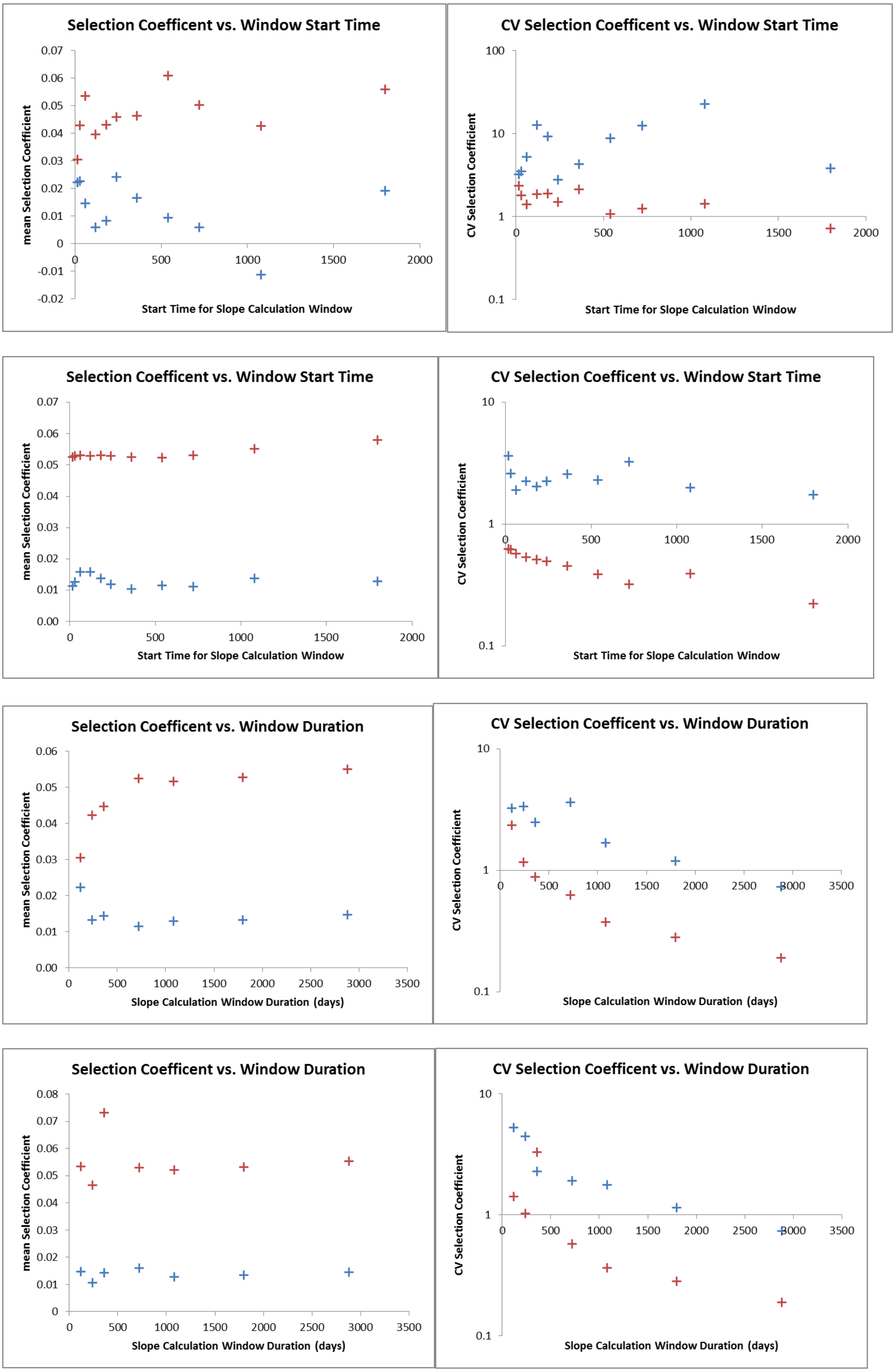
Selection Coefficent vs. Window Start Time
mean Selection Coefficient
0.07
0.06
0.05
0.04
0.03
0.02
0.01
0
-0.01
-0.02
0
500
1000
1500
2000
Start Time for Slope Calculation Window
CV Selection Coefficent vs. Window Start Time
CV Selection Coefficient
100
10
1
0.1
0
500
1000
1500
2000
Start Time for Slope Calculation Window
Selection Coefficent vs. Window Start Time
mean Selection Coefficient
0.07
0.06
0.05
0.04
0.03
0.02
0.01
0.00
0
500
1000
1500
2000
Start Time for Slope Calculation Window
CV Selection Coefficent vs. Window Start Time
CV Selection Coefficient
10
1
0.1
0
500
1000
1500
2000
Start Time for Slope Calculation Window
Selection Coefficent vs. Window Duration
mean Selection Coefficient
0.06
0.05
0.04
0.03
0.02
0.01
0.00
0
500
1000
1500
2000
2500
3000
3500
Slope Calculation Window Duration (days)
CV Selection Coefficent vs. Window Duration
CV Selection Coefficient
10
1
0.1
0
500
1000
1500
2000
2500
3000
3500
Slope Calculation Window Duration (days)
Selection Coefficent vs. Window Duration
mean Selection Coefficient
0.08
0.07
0.06
0.05
0.04
0.03
0.02
0.01
0
0
500
1000
1500
2000
2500
3000
3500
Slope Calculation Window Duration (days)
CV Selection Coefficent vs. Window Duration
CV Selection Coefficient
10
1
0.1
0
500
1000
1500
2000
2500
3000
3500
Slope Calculation Window Duration (days)

Citations.


1. Kay, K. and I.M. Hastings, *Measuring windows of selection for anti-malarial drug treatments.* Malaria Journal, 2015. **14**(1): p. 1-10.
2. Smith, T., et al., *Relationship between the entomologic inoculation rate and the force of infection for Plasmodium falciparum malaria.* The American Journal of Tropical Medicine and Hygiene, 2006. **75**(2_suppl): p. 11-18.
